# Genetic information insecurity as state of the art

**DOI:** 10.1101/2020.07.08.192666

**Authors:** Garrett J. Schumacher, Sterling Sawaya, Demetrius King, Aaron J. Hansen

## Abstract

Genetic information is being generated at an increasingly rapid pace, offering advances in science and medicine that are paralleled only by the threats and risk present within the responsible ecosystem. Human genetic information is identifiable and contains sensitive information, but genetic data security is only recently gaining attention. Genetic data is generated in an evolving and distributed cyber-physical ecosystem, with multiple systems that handle data and multiple partners that utilize the data. This paper defines security classifications of genetic information and discusses the threats, vulnerabilities, and risk found throughout the entire genetic information ecosystem. Laboratory security was found to be especially challenging, primarily due to devices and protocols that were not designed with security in mind. Likewise, other industry standards and best practices threaten the security of the ecosystem. A breach or exposure anywhere in the ecosystem can compromise sensitive information. Extensive development will be required to realize the potential of this emerging field while protecting the bioeconomy and all of its stakeholders.

## 1. INTRODUCTION

Genetic information contained in nucleic acids, such as deoxyribonucleic acid (DNA), has become ubiquitous in society, enabled primarily by rapid biotechnological development and drastic decreases in DNA sequencing and DNA synthesis costs (Berger and Schneck, 2019; Naveed et al., 2015). Innovation in these industries has far outpaced regulatory capacity and remained somewhat isolated from the information security and privacy domains. A single human whole genome sequence can cost hundreds to thousands of dollars per sample, and when amassed genetic information can be worth millions^1,2^. This positions genetic information systems as likely targets for cyber and physical attacks.

Human genetic information is identifiable (Erlich et al., 2018; Lowrence and Collins, 2007) and also contains sensitive health information; yet it is not always defined in these capacities by law. Unlike most other forms of data, it is immutable, remaining with an individual for their entire life. Sensitive human genetic data necessitates protection for the sake of individuals, their relatives, and ethnic groups; genetic information in general must be protected to prevent national and global threats (Sawaya et al., 2020). Therefore, human genetic information is a uniquely confidential form of data that requires increased security controls and scrutiny. Furthermore, non-human biological sources of genetic data are also sensitive. For example, microbial genetic data can be used to create designer microbes with CRISPR-Cas and other synthetic biology techniques (Werner, 2019), presenting global and national security concerns.

Several genomics stakeholders have reported security incidents according to news sources^3,4,5,6^ and breach notifications^7,8,9,10,11,12^. The most common reasons were misconfigurations of cloud security settings and email phishing attacks, and one resulted from a stolen personal computer containing sensitive information^12^. The National Health Service’s Genomics England database in the United Kingdom has been targeted by malicious nation-state actors^13^, and 23andMe’s Chief Security Officer said their database of around ten million individuals is of extreme value and therefore “certainly is of interest to nation states”^14^. Despite this recognition, proper measures to protect genetic information are often lacking under current best practices in relevant industries and stakeholders. Multi-stakeholder involvement and improved understanding of the security risks to biotechnology are required in order to develop appropriate countermeasures (Millett et al., 2019). Towards these goals, this paper expands upon a microbiological genetic information system assessment by Fayans et al. (Fayans et al., 2020) to include a broader range of genetic information, as well as novel concepts and additional threats to the ecosystem.

## 2. INFORMATION SECURITY CONCEPTS FOR GENETICS

Confidentiality, integrity, and availability are the core principles governing the secure operation of a system (Fayans et al., 2020; International Organization for Standardization, 2012). Confidentiality is the principle of ensuring access to information is restricted based upon the information’s sensitivity. Examples of confidentiality include encryption, access controls, and authorization. Integrity is the concept of protecting information from unauthorized modification or deletion, while availability ensures information is accessible to authorized parties at all times. Integrity examples include logging events, backups, minimizing material degradation, and authenticity verification. Availability can be described as minimizing the chance of data or material destruction, as well as network, power, and other infrastructure outages. Sensitive genetic information, which includes both biological material and digital genetic data, is the primary asset of concern, and associated assets, such as metadata, electronic health records and intellectual property, are also vulnerable within this ecosystem.

### 2.1 Genetic information security classifications

Genetic information can be classified into two primary levels, sensitive and nonsensitive, based upon value, confidentiality requirements, criticality, and inherent risk. Sensitive genetic information can be further categorized into restricted and private sublevels.

❖ *Restricted Sensitive Genetic Information* can be expected to cause significant risk to a nation, ethnic group, individual, or stakeholder if it is disclosed, modified, or destroyed without authorization. The highest level of security controls should be applied to restricted sensitive genetic information. Examples of restricted sensitive information are material and data sourced from humans, resources humans rely upon, and organisms and microbes that could cause harm to humans or resources humans rely upon. Due to its identifiability, human genetic information can be especially sensitive and thus requires special security considerations.
❖ *Private Sensitive Genetic Information* can be expected to cause a moderate level of risk to a nation, ethnic group, individual, or stakeholder if it is disclosed, modified, or destroyed without authorization. Genetic information that is not explicitly classified as restricted sensitive or nonsensitive should be treated as private sensitive information. A reasonable level of security controls should be applied to private sensitive information. Examples of private sensitive information are intellectual property from research, breeding, and agricultural programs.
❖ *Nonsensitive (or Public) Genetic Information* can be expected to cause little risk if it is disclosed, modified, or destroyed without authorization. While few controls may be required to protect the confidentiality of nonsensitive genetic information, controls should be in place to prevent unauthorized modification or destruction of nonsensitive information. Examples of nonsensitive information are material and data sourced from non-human entities that are excluded from the sensitive level if the resulting data are to be made publicly available within reason.

### 2.2 Motivation and necessity for securing genetic information

The genetic information ecosystem can be compromised in numerous ways, including purposefully adversarial activities and human error. Organizations take steps to monitor and prevent error, and molecular biologists are skilled in laboratory techniques; however, they commonly do not have the expertise and resources to securely configure and operate these environments, nor are they enabled to do so by vendor service contracts and documentation. Basic security features and tools, such as antivirus software, can easily be subverted, and advanced protections are not commonly implemented. Much genetic data is already publicly available via open and semi-open databases, and dissemination practices are not properly addressed by regulations. There are wide-ranging motives behind adversaries targeting non-public genetic information (Fayans et al., 2020). Numerous stakeholders, personnel, and insecure devices are relied upon from the path of sample collection to data dissemination. Depending on the scale of an exploit, hundreds to millions of people could be compromised. Local attacks could lead to certain devices, stakeholders, and individuals being affected, while supply chain and remote attacks could lead to global-scale impact.

Widespread public dissemination and lack of inherent security controls equate to millions of individuals and their relatives having substantial risk imposed upon them. Genetic data can be used to identify an individual (Lin et al., 2004) and predict their physical characteristics (Li et al., 2019; Lippert et al., 2017), and capabilities for familial matching are increasing, with the ability to match individuals to distant relatives (Erlich et al., 2018; Edge et al., 2017; Ney et al., 2018). Identifiability of genetic information is a critical challenge leading to growing consumer privacy concerns (Baig et al., 2020), and behavioral predictions from genetic information are gaining traction to produce stronger predictors year over year (Gard et al., 2019; Johnson et al., 2008). Furthermore, many diseases and negative health outcomes have genetic determinants, meaning that genetic data can reveal sensitive health information about individuals and families (Sawaya et al., 2020).

These issues pale in comparison to the weaponization of genetic information. Genetics can inform both a doctor and an adversary in the same way, revealing weaknesses that can be used for treatment or exploited to cause disease (Sawaya et al., 2020). The creation of bioweapons utilizes the same processes as designing vaccines and medicines to mitigate infectious diseases, namely access to an original infectious organism or microbe and its genetic information (Berger and Roderick, 2014). This alarming scenario was thought to be unlikely only six years ago as the necessary specialized skills and expertise were not widely distributed. Since then, access to sensitive genetic data has increased, such as the genome sequences of the 2019 novel coronavirus (SARS-CoV-2) (Sah et al., 2020)^15^, African swine fever (Mazur-Panasiuk et al., 2019), and the 1918 Spanish influenza A (H1N1) (Tumpey et al., 2005) viruses. Synthetic biology capabilities, skill sets, and resources have also proliferated (Ney et al., 2017). SARS-CoV-2 viral clone creation from synthetic DNA fragments was possible only weeks after the sequences became publicly available (Thao et al., 2020). This same technology can be utilized to modify noninfectious microbes and microorganisms to create weaponizable infectious agents (Berger and Roderick, 2014; Chosewood and Wilson, 2009; Salerno and Koelm, 2002). COVID-19 susceptibility, symptoms, and mortality all have genetic components (Taylor et al., 2020; Ellinghaus et al., 2020; Nguyen et al., 2020), demonstrating how important it will be to safeguard genetic information in the future to avoid targeted biological weapons. Additionally, microbiological data cannot be determined to have infectious origins until widespread infection occurs or until it is sequenced and deeply analyzed (Chosewood and Wilson, 2009; Salerno and Koelm, 2002); hence, data that is potentially sensitive also needs to be protected throughout the entire ecosystem.

## 3. THE GENETIC INFORMATION THREAT LANDSCAPE

The genetic information ecosystem is a distributed cyber-physical system containing numerous stakeholders (Supplementary Material, Appendix 1), personnel, and devices for computing and networking purposes. The ecosystem is divided into the pre-analytical, analytical, and post-analytical phases that are synonymous with: (i) collection, storage, and distribution of biological samples, (ii) generation and processing of genetic data, and (iii) storage and sharing of genetic data (Supplementary Material, Appendix 2). This ecosystem introduces many pathways, or attack vectors, for malicious access to information and systems (Figure 1).

**Figure 1.**
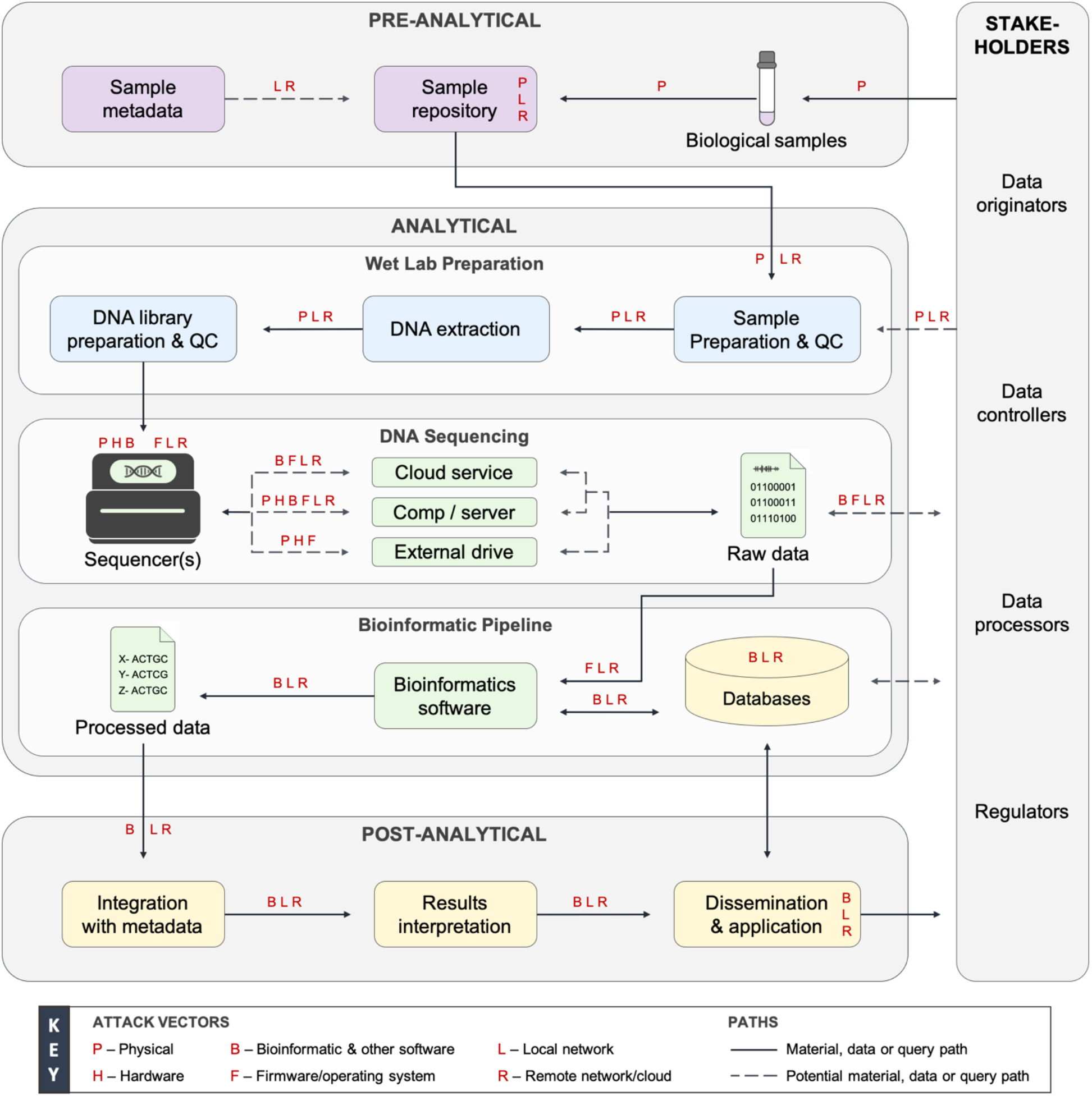
The genetic information ecosystem and accompanying threat landscape. The genetic information ecosystem is divided into three phases: pre-analytical, analytical, and post-analytical. The analytical phase is further divided into wet laboratory preparation, DNA sequencing, and bioinformatic pipeline subphases. In its simplest form, this system is a series of inputs and outputs that are either biological material, data, or logical queries on data. Every input, output, device, process, and stakeholder are vulnerable to exploitation via the attack vectors denoted by red letters. Color schema: purple, sample collection and processing; blue, wet laboratory preparation; green, genetic data generation and processing; yellow, data dissemination, storage, and application. Figure modified from Fayans et al. 2020 with permissions. More information on the ecosystem is provided in Supplementary Material, Appendix 2. Acronyms: QC, quality control; Comp, computing device.

### 3.1 Personnel and physical access controls

Unauthorized physical access or insider threats could allow for theft of assets or the use of other attack vectors on any phase of the ecosystem (Walsh and Streilein, 2020). Small independent laboratories do not often have resources to implement strong physical security. Large institutions are often enabled to maintain strong physical security, but the relatively large number of individuals and devices that need to be secured can create a complex attack surface. Ultimately, the strongest cybersecurity can be easily circumvented by weak physical security. Insider threats are a problem for information security because personnel possess deeper knowledge of an organization and its systems. Many countries rely on foreign nationals working in biotechnological fields that may be susceptible to foreign influence^16^. Citizens can also be susceptible to foreign influence^17^. Personnel could introduce many exploits on-site if coerced or threatened. Even when not acting in a purposefully malicious manner, personnel can unintentionally compromise the integrity and availability of genetic information through error (US Office of the Inspector General, 2004). Appropriate safeguards should be in place to ensure that privileged individuals are empowered to do their work correctly and efficiently, but all activities should be documented and monitored when working with sensitive genetic information.

### 3.2 Biological samples, metadata and repositories

Sample collection, storage, and distribution processes have received little recognition as legitimate points for the compromise of genetic information. Biological samples as inputs into this ecosystem can be modified maliciously to contain encoded malware (Ney et al., 2017), or they could be degraded, modified, or destroyed to compromise the material’s and resulting data’s integrity and availability. Sample repository and storage equipment are usually connected to a local network for monitoring purposes. A remote or local network attack could sabotage connected storage equipment, causing samples to degrade or be destroyed. Biorepositories and the collection and distribution of samples could be targeted to steal numerous biological samples, such as in known genetic testing scams^18^. Targeted exfiltration of small numbers of samples may be difficult to detect. Sensitive biological material should be safeguarded in storage and transit, and when not needed for long-term biobanking, it should be destroyed following successful analysis. Other organizations that handle genetic material could be targeted for the theft of samples and processed DNA libraries. The wet laboratory preparation and DNA sequencing subphases last several weeks and produce unused waste and stored material. At the conclusion of sequencing runs, the consumables that contain DNA molecules are not always considered sensitive. These items can be found unwittingly maintained in many sequencing laboratories. Several cases have been documented of DNA being recovered and successfully sequenced while aged for years at room temperature and in non-controlled environments (Colette et al., 2011).

### 3.3 Laboratories and equipment

DNA sequencing systems and laboratories are multifaceted in their design and threat profile. DNA sequencing instruments have varying scalability of throughput, cost, and unique considerations for secure operation (Table 1). Sequencing instruments have a built-in computer and commonly have connected computers and servers for data storage, networking, and analytics. These devices contain a number of different hardware components, firmware, an operating system, and other software. Some contain insecure legacy versions of operating system distributions. Sequencing systems usually have wireless or wired local network connections to the Internet that are required for device monitoring, maintenance, data transmission, and analytics in most operations. Wireless capabilities and Bluetooth technology within laboratories present unnecessary threats to these systems, as any equipment connected to laboratory networks is a potential network entry point.

**Table 1.**
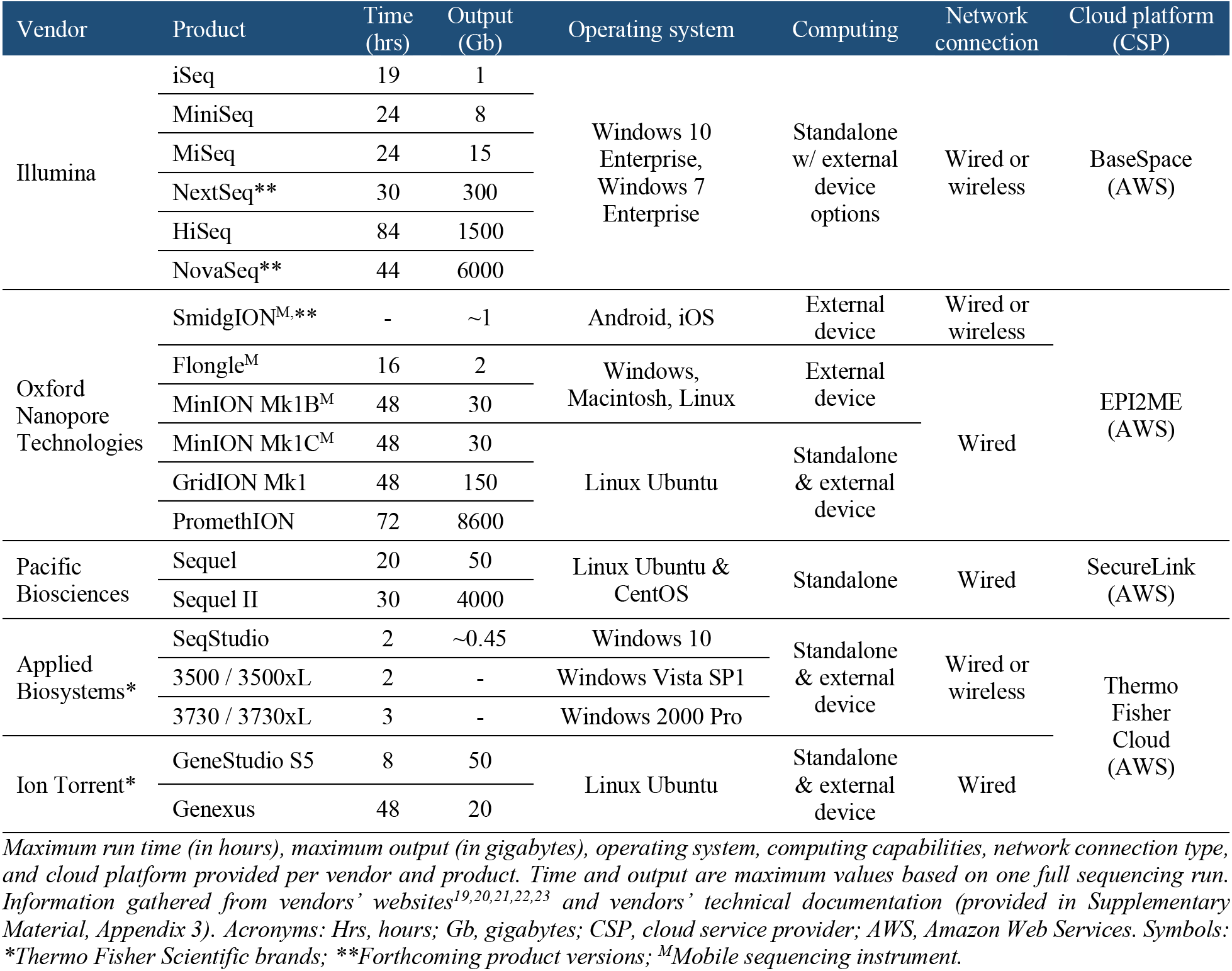
Overview of genetic sequencing systems.

Device vendors obtain various internal hardware components from several sources and integrate them into laboratory devices that contain vendor-specific intellectual property and software. Generic hardware components are often produced overseas, which is cost effective but leads to insecurities and a lack of hardening for specific end-use purposes. Hardware vulnerabilities could be exploited on-site, or they can be implanted during manufacturing and supply-chain processes for widespread and unknown security issues (Fayans et al., 2020; Ender et al., 2020; Shwartz et al., 2017; Anderson and Kuhn, 1997). Such hardware issues are unpatchable and will remain with devices forever until newer devices can be manufactured to replace older versions. Unfortunately, adversaries can always shift their techniques to create novel vulnerabilities within new hardware in a continual vicious cycle.

Third-party manufacturers and device vendors implement firmware in these hardware components. Embedded device firmware has been shown to be more susceptible to cyber-attacks than other forms of software (Shwartz et al., 2017). In-field upgrades are difficult to implement, and like hardware, firmware and operating systems of sequencing systems can be maliciously altered within the supply chain (Fayans et al., 2020). A firmware-level exploit would allow for the evasion of operating system, software, and application-level security features. Firmware exploits can remain hidden for long periods, even after hardware replacements or wiping and restoring to default factory settings. Furthermore, operating systems have specific disclosed Common Vulnerabilities and Exposures (CVEs) that are curated by the MITRE organization and backed by the US government^24^. With ubiquitous implementation in devices across all phases of the ecosystem, these software issues are especially concerning but can be partially mitigated by frequent updates. However, operating systems and firmware are typically updated every six to twelve months by a field agent accessing a sequencing device on site. Device operators are not allowed to modify the device in any way, yet they are responsible for some security aspects of this equipment. Additionally, researchers have confirmed the possibility of index hopping, or index misassignment, by sequencing device software, resulting in customers receiving confidential data from other customers (Ney et al., 2017) or downstream data processors inputting incorrect data into their analyses.

DNA sequencing infrastructure is proliferating. Illumina, the largest vendor of DNA sequencing instruments, accounted for 90% of the world’s sequencing data in 2017 by their own account^20^. In 2018, Illumina had 9,970 sequencers implemented globally capable of producing a total daily output of 893 Tb (Erlich, 2018), with many of these instruments housed outside of the US and Europe. In 2020, technology developed by Beijing Genomics Institute has finally resulted in the $100 human genome (Drmanac, 2020) while US prices remain around $1,000. Overseas organizations can be third-party sequencing service providers for direct-to-consumer (DTC) companies and other stakeholders. Shipping internationally for analysis is less expensive than local services (Office of the US Trade Representative, 2018), indicating that genetic data could be aggregated globally by nation-states^25^ and other actors during the analysis phase.

### 3.5 Storage and compute infrastructure

Raw signal sequencing data are stored on a sequencing system’s local memory and are transmitted to one or more endpoints. Transmitting data across a local network requires internal information technology (IT) configurations. Vendor documentation usually depends upon implementing a firewall to secure sequencing systems, but doing so correctly requires deep knowledge of secure networking and vigilance of network activity. Documentation also commonly mentions disabling and enabling certain network protocols and ports and further measures that can be difficult for most small-to medium-sized organizations if they lack dedicated IT support.

Laboratories and DNA sequencing systems are connected to many third-party services, and laboratories have little control over the security posture of these connections. Independent cloud platforms and DNA sequencing vendors’ cloud platforms are implemented for bioinformatic processing, data storage, and device monitoring and maintenance capabilities (Table 1). A thorough security assessment of cloud services remains unfulfilled in the genomics context. Multifactor authentication, role- and task-based access, and many other security measures are not common in these platforms. Misconfigurations to cloud services and remote communications are a primary vulnerability to genetic information, demonstrated by prior breaches, Remote Desktop Protocol issues affecting Illumina devices^26^, and a disclosed vulnerability in Illumina’s BaseSpace application program interface^27^. Laboratory information management systems (LIMS) are also frequently implemented within laboratories and connected to sequencing systems and laboratory networks (Roy et al., 2016). DNA sequencing vendors provide their own LIMSs as part of their cloud offerings. Even when LIMS and cloud platforms meet all regulatory requirements for data security and privacy, they are handling data that is not truly anonymized and therefore remains identifiable and sensitive. Furthermore, specific CVEs have been disclosed for dnaTools’ dnaLIMS product^28^ that were actively exploited by a foreign nation-state^29^. Phishing attacks are another major threat, as email services add to the attack surface in many ways. Sequencing service providers often share links granting access to datasets via email. These email chains are a primary trail of transactions that could be exploited to exfiltrate data on clients, metadata of samples, or genetic data itself.

Some laboratories transmit raw data directly to an external hard drive per customer or regulatory requirements. Reducing network activity in this way can greatly minimize the threat surface of sensitive genetic information. Separating networks and devices from other networks, or air gapping, while using hard drives is possible, but even air-gapped systems have been shown to be vulnerable to compromise (Guri, 2020; Guri et al., 2019). Sequencing devices are still required to be connected to the Internet for maintenance and are often connected between offline operations. Hard drives can be physically secured and transported; however, these methods are time and resource intensive, and external drives could be compromised for the injection of modified software or malware.

### 3.6 Bioinformatic pipeline

Bioinformatic software has not been commonly scrutinized in security contexts or subjected to the same adversarial pressure as other more mature software. Open-source software is widely used across genomics, acquired from several online code repositories, and heavily modified for individual purposes, but it is only secure when security researchers are incentivized to assess these products. In a specialized and niche industry like genomics and bioinformatics this is typically not the case. Bioinformatic programs have been found to be vulnerable due to poor coding practices, insecure function usage, and buffer overflows^30,31^ (Ney et al., 2017). Many researchers have uncovered that algorithms can be forced to mis-classify by intentionally modifying data inputs, breaking the integrity of any resulting outputs (Finlayson et al., 2019). Nearly every imaginable algorithm, model type, and use case has been shown to be vulnerable to this kind of attack across many data types, especially those relevant to raw signal and sequencing data formats (Biggio and Roli, 2018). Similar attacks could be carried out in the processing of raw signal data internal to a sequencing system or on downstream bioinformatic analyses accepting raw sequencing data or processed data as an input.

### 3.7 Dissemination practices and databases

Alarming amounts of human and other sensitive genetic data are publicly available^32,33,34,35,36^. Several funding and publication agencies require public dissemination, so researchers commonly contribute to open and semi-open databases (Shi and Wu, 2017). Healthcare providers either house their own internal databases or disseminate to third-party databases. Their clinical data is protected like any other healthcare information as required by regulations; however, this data can be sold and aggregated by external entities. DTC companies keep their own internal databases closely guarded and can charge steep prices for third-party access. Data sharing is prevalent when the price is right. Data originators often have access to their genetic data and test results for download in plaintext. These reports can then be uploaded to public databases, such as GEDmatch and DNA.Land, for further analyses, including finding distant genetic relatives with a shared ancestor (Erlich et al., 2018). A well-known use of such identification tactics was the infamous Golden State Killer case (Edge and Coop, 2019). Data sharing is dependent upon the data controller’s wants and needs, barring any legal or business requirements from other involved stakeholders. Genetic database vulnerabilities have been well-studied and disclosed (Edge and Coop, 2020; Ney et al., 2020; Naveed et al., 2015; Erlich and Narayanan, 2014; Gymrek et al., 2013). For example, the contents of the entire GEDmatch database could be leaked by uploading artificial genomes (Ney et al., 2020). Such an attack would violate the confidentiality of more than a million users’ and their relatives’ genetic data because the information is not truly anonymized. Even social media posts can be filtered for keywords indicative of participation in genetic research studies to identify research participants in public databases (Liu et al., 2019). All told, tens of millions of research participants, consumers, and relatives are already at risk.

### 3.8 Perceived risk to phases of the genetic information ecosystem

Adversarial targeting of genetic information largely depends upon the sensitivity, quantity, and efficiency of information compromise for attackers, leading to various states in likelihood of a breach or exposure scenario. The impact of a compromise is determined by a range of factors, including the size of the population at risk, negative consequences to stakeholders, and capabilities and scale of adversarial activity. Likelihood and impact both ultimately inform the level of risk facing stakeholders during ecosystem phases (Figure 2).

**Figure 2.**
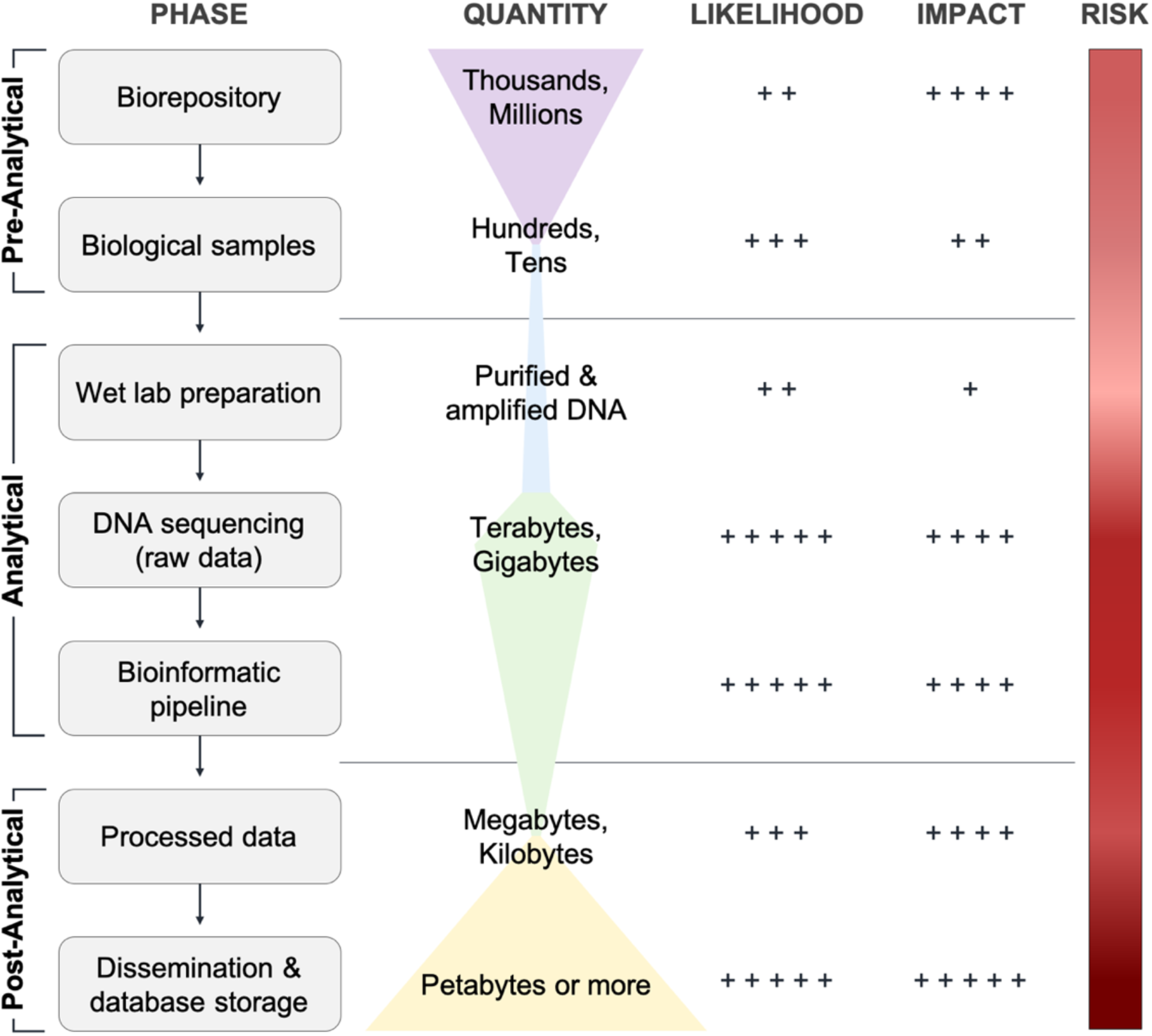
Risk to the genetic information ecosystem. Quantity is not to scale but is denoted abstractly by width of the second column. Likelihood judged by the available threats and opportunities to adversaries and the efficiency of an attack. Impact in terms of the number of people affected and the current and emerging consequences to stakeholders. Likelihood and impact scores: Low (+); Moderate (+ +); High (+ + +); Very High (+ + + +); Extreme (+ + + + +). Low to extreme risk is denoted by the hue of red, from light to dark.

## 4. DISCUSSION

Security is a spectrum; stakeholders must do everything they can to chase security as a best practice. Securing genetic information is a major challenge in this rapidly evolving ecosystem. Attention has primarily been placed on the post-analytical phase of the genetic information ecosystem for security and privacy, but adequate measures have yet to be universally adopted. The pre-analytical and analytical phases are also highly vulnerable points for data compromise that must be addressed. Adequate national regulations are needed for security and privacy enforcement, incentivization, and liability, but legal protection is dictated by regulators’ responses and timelines. However, data originators, controllers, and processors can take immediate action to protect their data.

Genetic information security is a shared responsibility between sequencing laboratories and device vendors, as well as all other involved stakeholders. To protect genetic information, laboratories, biorepositories, and other data processors need to create strong organizational policies and reinvestments towards their physical and cyber infrastructure. They also need to determine the sensitivity of their data and material and take necessary precautions to safeguard sensitive genetic information. Data controllers, especially healthcare providers and DTC companies, should reevaluate their data sharing models and methods, with special consideration for the identifiability of genetic data. Device vendors need to consider security when their products are being designed and manufactured. Many of these recommendations go against the current paradigms in genetics and related industries and will therefore take time, motivation, and incentivization before being actualized, with regulation being a critical factor. In order to secure genetic information and protect all stakeholders in the genetic information ecosystem, further in-depth assessments of this threat surface will be required, and novel security and privacy approaches will need to be developed. Sequencing systems, bioinformatics software, and other biotechnological infrastructure need to be analyzed to fully understand their vulnerabilities. This will require collaborative engagement between stakeholders to implement improved security measures into genetic information systems (Moritz et al., 2020; Berger and Schneck, 2019). The development and implementation of genetic information security will foster a healthy and sustainable bioeconomy without damaging privacy or security.

There can be security without privacy, but privacy requires security. These two can be at odds with one another in certain contexts. For example, personal security aligns with personal privacy, whereas public security can require encroachment on personal privacy. A similar story is unfolding within genetics. Genetic data must be shared for public good, but this can jeopardize personal privacy. However, genetic data necessitates the strongest protections possible for public security and personal security. Appropriate genetic information security will simultaneously protect everyone’s safety, health, *and* privacy.

## 5. METHODOLOGY

The inspiration for this work occurred while performing several security assessments and penetration tests of DNA sequencing laboratories and other stakeholders. Initially, an analysis of available literature and technical documentation (n=57) was performed, followed by confidential semi-structured interviews (n=46) with key personnel from multiple relevant stakeholders. The study’s population consisted of leaders and technicians from government agencies (n=3) and organizations in small, medium, and large enterprises (n=18) across the United States, including California, Colorado, District of Columbia, Massachusetts, Montana, and Virginia. Several stakeholders allowed access to their facilities for observing environments and further discussions. Some stakeholders allowed in-depth assessments of equipment, networks, and services.

## CONFLICTS OF INTEREST

GS, SS, and DN are founders and owners of GeneInfoSec Inc. and are developing technology and services to protect genetic information. GeneInfoSec Inc. has not received US Federal research funding.

AH declares that the research was conducted in the absence of any commercial or financial relationships that could be construed as a potential conflict of interest.

## AUTHOR CONTRIBUTIONS

Inception - GS. Literature review and analysis - GS, SS, DN. Stakeholder engagement and interviews - GS, SS, DN, AH. Security assessments - GS, AH. Drafting of paper - GS. Critical review of draft - GS, SS, DN, AH.

## ACKNOWLEDGEMENTS

The authors would like to acknowledge the confidential research participants and collaborators on this study for their time, resources, and interest in bettering genetic information security. Thank you to Cory Cranford, Arya Thaker, Ashish Yadav, and Dr. Kevin Gifford and Dr. Daniel Massey of the Department of Computer Science, formerly of the Technology, Cybersecurity and Policy Program, at the University of Colorado Boulder for their support of this work.

## SUPPLEMENTARY MATERIAL

Accompanying appendices of supplementary material are attached to the end of this manuscript. Material provided includes:

❖ Appendix 1. Overview and examples of genetic information ecosystem stakeholders, and Supplementary Table 1 (page 19)
❖ Appendix 2. Overview of the genetic information ecosystem processes (page 21)
❖ Appendix 3. Genetic sequencing system vendor documentation (page 23)

## SUPPLEMENTARY MATERIAL

## Appendix 1. Overview and examples of genetic information ecosystem stakeholders

Genetics stakeholders are categorized based upon their influence, contributions, and handling of biological samples and resulting genetic data (Supplementary Table 1). Asymmetries exist between stakeholders in these regards^37^. *Data originators* are humans that voluntarily or involuntarily are the source of biological samples or are investigators collecting samples from nonhuman specimens. Examples of data originators include consumers, healthcare patients, military personnel, research subjects, migrants, criminals, and their relatives. *Data controllers* are entities that are legally liable for and dictate the use of biological samples and resulting data. In human-derived contexts, data controllers are typically healthcare providers, researchers, law enforcement agencies, or DTC companies. *Data processors* are entities that collect, store, generate, analyze, disseminate, and/or apply biological samples or genetic data. Data processors may also be data originators and data controllers. Examples include biorepositories, DNA sequencing laboratories, researchers, cloud and other service providers, and supply chain entities responsible for devices, software and materials. *Regulators* oversee this ecosystem and the application and use of biotechnology, biological samples, genetic data, and market/industry trends at the transnational, national, local, and organizational levels.

**Supplementary Table 1.** Categories and groups of genetic information ecosystem stakeholders with examples of organizations and contexts.

| Stakeholder | Group | Examples / context |
| --- | --- | --- |
| Data originators | Consumers, patients, subjects | Humans voluntarily providing biological samples |
|  | Researchers | Investigators collecting non-human samples |
|  | Suspected criminals, migrants, relatives | Humans involuntarily providing biological samples |
| Data controllers | Data originators | Rare instances of individual data control, e.g., Genetic Alliance |
|  | DTC companies | 23andme, Ancestry.com, FamilyTreeDNA, MyHeritage |
|  | Healthcare providers | Kaiser, UnitedHealth, NorthShore, Veterans Affairs (VA) |
|  | Research organizations | Academic, military, government and industry groups |
| Data processors | Biorepositories / biobanks | All of Us (NIH), UK Biobank, Beryllium BioBank |
|  | Cloud service providers | AWS, Google Cloud, Box, LifeOmic, BioBright, BlueBee |
|  | Data controllers | DTC, healthcare & research groups processing data |
|  | Device manufacturers/vendors | Microsoft, Apple, Dell, Linux-based groups, Agilent, Qiagen |
|  | Diagnostics providers | Quest Diagnostics, Ambry Genetics, academic laboratories |
|  | Email service providers | Gmail, Microsoft Outlook, ProtonMail |
|  | IT organizations | Internal and third-party IT services, Internet providers |
|  | Open and semi-open databases | GEDmatch, All of Us (NIH), Genomics England (NHS) |
|  | Sequencing laboratories | Third-party and internal sequencing service providers |
|  | Sequencing system vendors | Illumina, Oxford Nanopore, Thermo Fisher, Pacific Biosciences |
|  | Synthetic DNA providers | Ginkgo, IDT, Agilent, Twist Biosciences, Thermo Fisher |
| Regulators | Consumers | Dictate aspects of ecosystem through behavior |
|  | Federal organizations | US Centers for Medicare & Medicaid Services, FDA |
|  | International and national governments | United States, European Union, United Kingdom |
|  | Intranational governments | Providences, states, counties, territories, tribes |
|  | Law enforcement | FBI, DHS, DOJ, military, police |
|  | Organizational policy creators | Management and executive personnel |

*Acronyms: DTC, direct-to-consumer; NIH, US National Institutes of Health; UK, United Kingdom; IT, information technology; NHS, UK National Health Services; IDT, Integrated DNA Technologies; US, United States; FDA, US Food and Drug Administration; FBI, US Federal Bureau of Investigation; DHS, US Department of Homeland Security; DOJ, US Department of Justice.*

## Appendix 2. Overview of the genetic information ecosystem processes

Biological samples and metadata from the samples must first be collected once a data originator or controller determines to proceed with genetic testing. Biological samples can be sourced from any biological entity relying on nucleic acids for reproduction, replication, and other processes, including non-living microbes (e.g., viruses, prions), microorganisms (e.g., bacteria, fungi), and organisms (e.g., plants, animals). Samples are typically de-identified of metadata and given a numeric identifier, but this is largely determined by the interests of data controllers and the regulations that may pertain to various sample types. Metadata includes demographic details, inclusion and exclusion criteria, pedigree structure, health conditions critical for secondary analysis, and other identifying information^38^. It can also be in the form of quality metrics obtained during the analysis phase. Samples are then stored in controlled environments at decreased temperature, moisture, light, and oxygen to avoid degradation. Sample repositories can be internal or third-party infrastructure housing small to extremely large quantities of material for short- and long-term storage. Following storage, samples are distributed to an internal or third-party laboratory for DNA sequencing preparations.

The wet laboratory preparation phase chemically prepares biological samples for sequencing with sequencing-platform-dependent methods. This phase can be performed manually with time- and labor-intensive methods, or it can be highly automated to reduce costs, run-time, and error. Common initial preparation steps involve removing contaminants and unwanted material from biological samples and extracting and purifying samples’ nucleic acids. If RNA is to be sequenced, it is usually converted into complementary DNA. Once DNA has been isolated, a library for sequencing is created via size-selection, sequencing adapter ligation, and other chemical processes. Adapters are synthetic DNA molecules attached to DNA fragments for sequencing and contain sample indexes, or identifiers. Indexes allow for multiplexing sequencing runs with many samples at once to increase throughput, decrease costs, and to identify DNA fragments to their sample source.

To begin sequencing, prepared libraries are loaded into a DNA sequencing instrument with the required materials and reagents. Laboratory personnel must login to the instrument and any connected services, such as cloud services or information management systems, and configure a run to initiate sequencing. A single sequencing run can generate gigabytes to terabytes of raw sequencing data and last anywhere from a few hours to multiple days, requiring the devices to commonly be left unmonitored during operation. Raw data can be stored on the instrument’s local memory and are transmitted to one or more of the following endpoints during or following a sequencing run: (i) local servers, computers, or other devices within the laboratory; (ii) cloud services of the vendor or other service providers; and (iii) external hard drives directly tethered to the sequencer. Data paths largely depend on the sequencing platform, the laboratory’s capabilities and infrastructure, and the sensitivity of data being processed. Certain regulations require external hard drive use and offline data storage, analysis, and transmission.

Bioinformatic pipelines convert raw data through a series of software tools into usable forms. Raw signal data include images, chemical signal, electrical current signal, and other forms of signal data dependent upon the sequencing platform. Primary analyses convert raw signal data into sequence data with accompanying quality metrics through a process known as basecalling. Many sequencing instruments can perform these functions. The length of each DNA molecule sequenced is orders of magnitude smaller than genes or genomes of interest, so basecalled sequence data must then be aligned to determine each read’s position within a genome or genomic region. This aligned sequence data is then compared to reference genomes sourced from databases through a procedure known as variance detection to determine differences between a sample’s data and the accepted normal genomic sequence. Only the unique genetic variants of a sample are retained in variance call format (VCFs) files, a common final processed data form. VCF files are vastly smaller than the gigabytes to terabytes of raw data initially produced, making them an efficient format for longterm storage, dissemination, and analysis purposes. However, this file format exists as a security threat for sensitive genetic data because these files are personally identifiable and contain sensitive health information.

Following data analyses, processed data are integrated with metadata and ultimately interpreted for the data controller’s purpose. Metadata and genetic data are often housed together, and exploiting this combined information could lead to numerous risks and threats to the data originators, their relatives, and the liable entities involved along the data path. Secondary analyses can be performed on datasets by data controllers and third-party data processors to answer any number of relevant research questions, such as in diagnostics or ancestry analysis. Genetic research is only powerful when large datasets are created containing numerous data points from thousands to millions of samples. Therefore, genetic data is widely distributed and accessible via remote means across numerous databases and stakeholders.

## Appendix 3. Genetic sequencing system vendor documentation

Applied Biosystems, Life Technologies Corp. (Hitachi), Thermo Fisher Scientific, Inc. Applied Biosystems 3500/3500xL Genetic Analyzer User Guide. (2010). *Thermo Fisher Scientific publication #4401661, Rev. C*. Retrieved from http://tools.thermofisher.com/content/sfs/manuals/4401661.pdf.

Applied Biosystems, Life Technologies Corp. (Hitachi), Thermo Fisher Scientific, Inc. Applied Biosystems 3730/37300xL DNA Analyzers User Guide. (2010). *Thermo Fisher Scientific publication #444331468, Rev. E*. Retrieved from https://assets.thermofisher.com/TFS-Assets/LSG/manuals/cms_041259.pdf.

Applied Biosystems, Thermo Fisher Scientific, Inc. Applied Biosystems SeqStudio Genetic Analyzer Specification Sheet. (2020). *Thermo Fisher Scientific publication #COL23988 0320*. Retrieved from https://assets.thermofisher.com/TFS-Assets/GSD/Specification-Sheets/SeqStudio-Specification-Sheet.pdf.pdf.

Illumina, Inc. HiSeq X System Lab Setup and Prep Guide. (January 2017). *Illumina document #15050093 v05*. Retrieved from https://support.illumina.com/content/dam/illumina-support/documents/documentation/system_documentation/hiseqx/hiseq-x-lab-setup-and-site-prep-guide-15050093-05.pdf.

Illumina, Inc. Illumina Proactive | Data Security Sheet. (2019). *Illumina document #970-2019-019-A*. Retrieved from https://www.illumina.com/content/dam/illumina-marketing/documents/informatics/illumina-proactive-data-security-sheet-970-2019-019.pdf.

Illumina, Inc. iScan System Site Prep Guide. (January 2019). *Illumina document #1000000000661 v01*. Retrieved from https://support.illumina.com/content/dam/illumina-support/documents/documentation/system_documentation/iscan/iscan-system-site-prep-guide-1000000000661-01.pdf.

Illumina, Inc. iSeq 100 Sequencing System Site Prep Guide. (April 2020). *Illumina document #1000000035337 v07*. Retrieved from https://support.illumina.com/content/dam/illumina-support/documents/documentation/system_documentation/iseq100/iseq-100-site-prep-guide-1000000035337-07.pdf.

Illumina, Inc. MiniSeq System Guide. (February 2020). *Illumina document # 1000000002695 v03, Material #20014309*. Retrieved from https://support.illumina.com/content/dam/illumina-support/documents/documentation/system_documentation/miniseq/miniseq-system-guide-1000000002695-03.pdf.

Illumina, Inc. MiSeq System Guide. (August 2019). *Illumina document #15027617 v05, Material #20000262*. Retrieved from https://support.illumina.com/content/dam/illumina-support/documents/documentation/system_documentation/miseq/miseq-system-guide-for-local-run-manager-15027617-05.pdf.

Illumina, Inc. NextSeq 550Dx Instrument Site Prep Guide. (March 2019). *Illumina document #1000000009869 v03*. Retrieved from https://support.illumina.com/content/dam/illumina-support/documents/documentation/system_documentation/nextseq-550dx/nextseq-550dx-instrument-site-prep-guide-1000000009869-03.pdf.

Illumina, Inc. NovaSeq 6000 Sequencing System Site Prep Guide. (January 2019). *Illumina document #1000000019360 v06*. Retrieved from https://support.illumina.com/content/dam/illumina-support/documents/documentation/system_documentation/novaseq/novaseq-site-prep-guide-1000000019360-06.pdf.

Ion Torrent, Thermo Fisher Scientific, Inc.. Ion GeneStudio S5 Series Specification Sheet. (2018). *Thermo Fisher Scientific publication #COL22253 0118*. Retrieved from https://assets.thermofisher.com/TFS-Assets/CSD/Specification-Sheets/PG1720-PJT2769-COL22253-P-on-GeneStudio-S5-Spec-Sheet-Global-FLR.pdf.

Ion Torrent, Thermo Fisher Scientific, Inc.. Ion GeneStudio S5 Series Systems Brochure. (2018). *Thermo Fisher Scientific publication #COL06368 1018*. Retrieved from https://assets.thermofisher.com/TFS-Assets/CSD/brochures/ion-genestudio-s5-ngs-system-brochure.pdf.

Ion Torrent, Thermo Fisher Scientific, Inc.. Ion Torrent Genexus Integrated Sequencer Performance Summary Sheet. (2019). *Thermo Fisher Scientific publication #MAN0017918*. Retrieved from https://assets.thermofisher.com/TFS-Assets/LSG/manuals/MAN0017918_GenexusIntegratedSequencer_SPG.pdf.

Oxford Nanopore Technologies. GridION Mk1 Site Installation and Device Operation Requirements, version 1. (July 2019). Retrieved from https://nanoporetech.com/sites/default/files/s3/products/GridION%20Mk1%20IT%20Requirements_v1.pdf.

Oxford Nanopore Technologies. MinION IT requirements, version 3. (February 2020). Retrieved from https://community.nanoporetech.com/requirements_documents/minion-it-reqs.pdf.

Oxford Nanopore Technologies. PromethION P24/P48 Site Installation and Device Operation Requirements, version 1. (July 2019). Retrieved from https://nanoporetech.com/sites/default/files/s3/products/PromethION%20P24-P48%20IT%20Requirements_v1.pdf.

Pacific Biosciences of California, Inc. Operations Guide – Sequel System: The SMRT Sequencer. (October 2018). *PacBio publication #101-055-100-06*. Retrieved from https://www.pacb.com/wp-content/uploads/Operations-Guide-The-SMRT-Sequencer-Sequel-System.pdf.

Pacific Biosciences of California, Inc. Operations Guide – Sequel II System: The SMRT Sequencer. (April 2019). *PacBio publication #101-774-700-01*. Retrieved from https://www.pacb.com/wp-content/uploads/Operations-Guide-Sequel-II-System.pdf.

Thermo Fisher Scientific, Inc. Security Operations Guide: Connect Platform | IoT Connectivity, version 4.2. (2019). Retrieved from https://assets.thermofisher.com/TFS-Assets/CORP/Reference-Materials/Connect_whitepaper_Nov2019.pdf.

## Footnotes

1 https://www.gsk.com/en-gb/media/press-releases/gsk-and-23andme-sign-agreement-to-leverage-genetic-insights-for-the-development-of-novel-medicines/

2 https://www.ancestry.com/corporate/newsroom/press-releases/ancestrydna-and-calico-to-research-the-genetics-of-human-lifespan

3 https://www.bloomberg.com/news/articles/2019-11-06/breach-at-dna-test-firm-veritas-exposed-customer-information

4 https://www.bostonherald.com/2019/08/22/mgh-data-breach-exposes-10000-patients/

5 https://www.latimes.com/business/la-fi-vitagene-dna-privacy-exposed-20190709-story.html

6 https://www.komando.com/security-privacy/ancestry-com-suffers-big-data-leak-300000-user-credentials-exposed/435921/

7 https://www.ambrygen.com/legal/substitute-notice

8 https://media.dojmt.gov/wp-content/uploads/Data-Breach-NotificationDetails11.pdf

9 https://media.dojmt.gov/wp-content/uploads/Consumer-Notice-73.pdf

10 https://privacyrights.org/data-breaches/myriad-genetic-laboratories-inc

11 https://blog.myheritage.com/2018/06/myheritage-statement-about-a-cybersecurity-incident/

12 https://media.dojmt.gov/wp-content/uploads/Shire-Human-Genetic-Therapies-Inc.pdf

13 https://www.telegraph.co.uk/news/2018/12/05/nhs-storing-patients-genetic-data-high-security-army-base-due/

14 https://www.telegraph.co.uk/technology/2020/03/09/dna-testing-firms-risk-state-sponsored-hacks-says-23andme-security/

15 https://www.ncbi.nlm.nih.gov/genbank/sars-cov-2-seqs/

16 https://www.fbi.gov/investigate/counterintelligence/foreign-influence

17 https://www.sciencemag.org/news/2020/06/fifiy-four-scientists-have-lost-their-jobs-result-nih-probe-foreign-ties

18 https://oig.hhs.gov/fraud/consumer-alerts/alerts/geneticscam.asp

19 https://www.illumina.com/systems/sequencing-platforms.html

20 https://nanoporetech.com/products/

21 https://www.pacb.com/products-and-services/

22 https://www.thermofisher.com/us/en/home/brands/applied-biosystems.html

23 https://www.thermofisher.com/us/en/home/brands/ion-torrent.html

24 https://cve.mitre.org/

25 https://www.cisa.gov/news/2020/05/13/fbi-and-cisa-warn-against-chinese-targeting-covid-19-research-organizations

26 https://support.illumina.com/bulletins/2019/11/bluekeep-and-dejablue--two-vulnerabilities-of-the-remote-desktop.html

27 https://nvd.nist.gov/vuln/detail/CVE-2013-7111

28 https://www.shorebreaksecurity.com/blog/product-security-advisory-psa0002-dnalims/

29 https://www.zdnet.com/article/mysterious-iranian-group-is-hacking-into-dna-sequencers/

30 https://share-ng.sandia.gov/news/resources/news_releases/genomic_cybersecurity/

31 https://nvd.nist.gov/vuln/detail/CVE-2019-10269

32 https://my.pgp-hms.org/public_genetic_data

33 https://gnomad.broadinstitute.org/downloads

34 https://platform.stjude.cloud/data/diseases

35 https://www.ensembl.org/Homo_sapiens/Info/Index

36 https://www.completegenomics.com/public-data/

37 Berger, K. M., & Schneck, P. A. (2019). National and transnational security implications of asymmetric access to and use of biological data. *Frontiers in bioengineering and biotechnology, 7*, 21.

38 Erlich, Y., & Narayanan, A. (2014). Routes for breaching and protecting genetic privacy. *Nature Reviews Genetics, 15*(6), 409-421.

